# CRISPR/Cas9 mediated editing of the Quorn fungus *Fusarium venenatum* A3/5 by transient expression of Cas9 and sgRNAs targeting endogenous marker gene *PKS12*

**DOI:** 10.1101/2021.09.06.459129

**Authors:** Fiona M Wilson, Richard J Harrison

**Affiliations:** NIAB EMR, New Road, East Malling, West Malling, Kent ME19 6BJ; NIAB, 93 Lawrence Weaver Road, Cambridge CB3 0LE

**Keywords:** AMA1, CRISPR/Cas9, codon optimisation, *Fusarium venenatum*, PolII promoter, *5SrRNA*

## Abstract

**Background:** Gene editing using CRISPR/Cas9 is a widely used tool for precise gene modification, modulating gene expression and introducing novel proteins, and its use has been reported in a number of filamentous fungi including the genus *Fusarium*. The aim of this study was to optimise gene editing efficiency using AMA1 replicator vectors for transient expression of CRISPR constituents in *Fusarium venenatum* (A3/5), used commercially in the production of mycoprotein (Quorn™).

**Results:** We present evidence of CRISPR/Cas9 mediated gene editing in *Fusarium venenatum*, by targeting the endogenous visible marker gene *PKS12*, which encodes a polyketide synthase responsible for the synthesis of the pigment aurofusarin. Constructs for expression of single guide RNAs (sgRNAs) were cloned into an AMA1 replicator vector incorporating a construct for constitutive expression of *cas9* codon-optimised for *Aspergillus niger* or *F. venenatum*. Vectors were maintained under selection for transient expression of sgRNAs and *cas9* in transformed protoplasts. 100% gene editing efficiency of protoplast-derived isolates was obtained using *A. niger cas9* when sgRNA transcription was regulated by the *F. venenatum 5SrRNA* promoter. In comparison, expression of sgRNAs using a *PgdpA*-ribozyme construct was much less effective, generating mutant phenotypes in 0-40% of isolates, with evidence of off-target editing. Viable isolates were not obtained from protoplasts transformed with an AMA1 vector expressing *cas9* codon-optimised for *F. venenatum*.

**Conclusions:** Using an AMA1 replicator vector for transient expression of *A. niger cas9* and sgRNAs transcribed from the native *5SrRNA* promoter, we demonstrate efficient gene editing of an endogenous marker gene in *F. venenatum*, resulting in knockout of gene function and a visible mutant phenotype in 100% of isolates. This establishes a platform for further development of CRISPR/Cas technology in *F. venenatum*, such as modulation of gene expression, gene insertion, base editing and prime editing. These tools will facilitate an understanding of the controls of secondary metabolism and hyphal development during fermentation of *F. venenatum* for mycoprotein production and may be used to validate prototypes of strains for improvement using classical means, enabling more cost-effective and sustainable production of this industrially important fungus.

## Background

Quorn™ myco-protein is a high fibre, high quality protein with a significantly lower environmental impact compared to that of beef (1), produced by fermentation of *Fusarium venenatum* (ATCC 20334, A3/5) utilising wheat-derived glucose in a continuous fermentation process. The organism was selected for its efficient biomass conversion, nutritional value, sparsely branched mycelium which enables processing into a meat-like texture and absence of undesirable secondary metabolites under optimal growth conditions (2,3). *Fusarium venenatum* has no known sexual cycle and therefore strain improvement is only currently possible thorough clonal mutagenesis. There are multiple opportunities for strain improvement, including use of alternative carbon sources, which may require changes in strain response to be suitable for food production. These advances could be facilitated by biotechnological innovations including development of gene editing mediated by CRISPR/Cas9 as a research tool and potentially as an onward tool for strain improvement.

The aim of the current study was to optimise CRISPR/Cas9 mediated gene editing in *F. venenatum* and establish a platform for its use in understanding genetic controls of metabolic pathways and hyphal development. CRISPR/Cas9 is a well-established technology, successfully used in a wide range of organisms for precise gene editing, gene insertion and modulation, facilitating genomic functional analyses and production of novel compounds. The Cas9 enzyme, guided by a sequence-specific RNA complex (sgRNA), generates a double stranded break at the target site in the genome, which can be repaired by either the non-homologous end joining (NHEJ) or microhomology –mediated end joining mechanisms (which are error prone), or by homology-directed repair (HR) in the presence of a repair template.

CRISPR/Cas9 mediated gene editing in filamentous fungi was first reported in *Aspergillus* (4– 6) and *Trichoderma reesei* (7). Subsequent reports include use of CRISPR/Cas9 in *Penicillium chrysogenum* (8), *Ustilago maydis* (9) and *Magnaporthe oryzae* (10). Its application in *Fusarium* was reported initially in *F. graminearum* (11) and *F. oxysporum* (12) and subsequently in *F. proliferatum* (13) and *F. fujikuroi* (14).

Parameters for optimising the methodology in fungi have included use of various delivery systems for CRISPR constituents, evaluation of promoters driving sgRNA expression and codon optimisation of *cas9* and its fusion to appropriate protein nuclear localization signals (NLS).

Strategies used for delivery of CRISPR constituents include generation of stable transgenic lines for expression of *cas9* and sgRNAs (4,10,11), introduction of *in vitro* expressed sgRNAs (7,14,15), introduction of Cas9/sgRNA ribonucleic protein (RNP) complexes (8,10,12,13,15,16), transformation with expression vectors for transient production of Cas9 and sgRNAs *in vivo* (6,14), or sgRNAs produced *in vitro* (17). AMA1 replicator vectors for transient expression of sgRNAs and *cas9* have also been used successfully (5,8,9).

PolIII promoters are commonly used to drive sgRNA expression in fungi, such as the yeast SNR52 promoter (4,6) and endogenous promoters for *U6* (7–9,14,16), tRNA (8), *U3* (18) and *5SrRNA* (14,17). Others have expressed sgRNA embedded in a dual ribozyme driven by a PolII promoter (5,11).

Successful editing in fungi has been achieved using Cas9 proteins optimised for other organisms, such as human-optimized Cas9 (hSpCas9) (4,6,10,13,16) and *A. niger* optimised Cas9 for gene editing in *F. oxysporum* (12). Others report use of Cas9 optimised for the target organism (5,7,11,14).

We report on the use of an AMA1 vector system for transient expression of *cas9* and sgRNAs via protoplast transformation, for evaluating gene editing efficiency in *F. venenatum* of sgRNAs transcribed using the PolII *gdpA* promoter (*PgdpA*) within a dual ribozyme or by the *F. venenatum* PolIII *5SrRNA* promoter (*PFv5SrRNA*). We additionally compare gene editing efficiency of Cas9 optimised for *A. niger* (*A. niger* Cas9) or *F. venenatum* (*Fv* Cas9).

## Results

### Vector construction

For comparison of *Fv* Cas9 and *A. niger* Cas 9 a codon-optimised *FvCas9* gene was synthesised and an expression cassette *Ptef1-Fvcas9*-SV40-*Ttef1* was assembled (Additional file 1: Table S1) and ligated with the backbone of AMA1 vector pFC332 (5), replacing the *A. niger cas9* cassette to give vector pFCFvCas9. These vectors were used as ‘empty vector’ controls in transformation experiments. sgRNAs targeting two separate sites (numbered ‘14’ and ‘29’) in exon 3 of the endogenous *PKS12* gene were cloned downstream of promoter *PgdpA* within a dual ribozyme cassette (5) (giving constructs PolII-PK3/14 and PolII-PK3/29) or promoter *PFv5SrRNA* (giving constructs 5S-PK3/14 and 5S-PK3/29) (Figure 1, and see Additional file 2: Table S2 for *PFv5SrRNA* cassettes sequence). Promoter-sgRNA constructs were cloned into the USER site of pFC332 to give single sgRNA vectors pFC332::PolII-PK3/14 and pFC332::PK3/29. A dual PolII-sgRNA vector (pFC332::PolII-PK3/14/29) was also constructed. For comparison of gene editing efficiency with *A. niger cas9*, PolII-PK3/14 was additionally cloned into the USER site of vector pFCFvCas9 to give pFCFvCas9::PolII-PK3/14,.

**Fig. 1.**
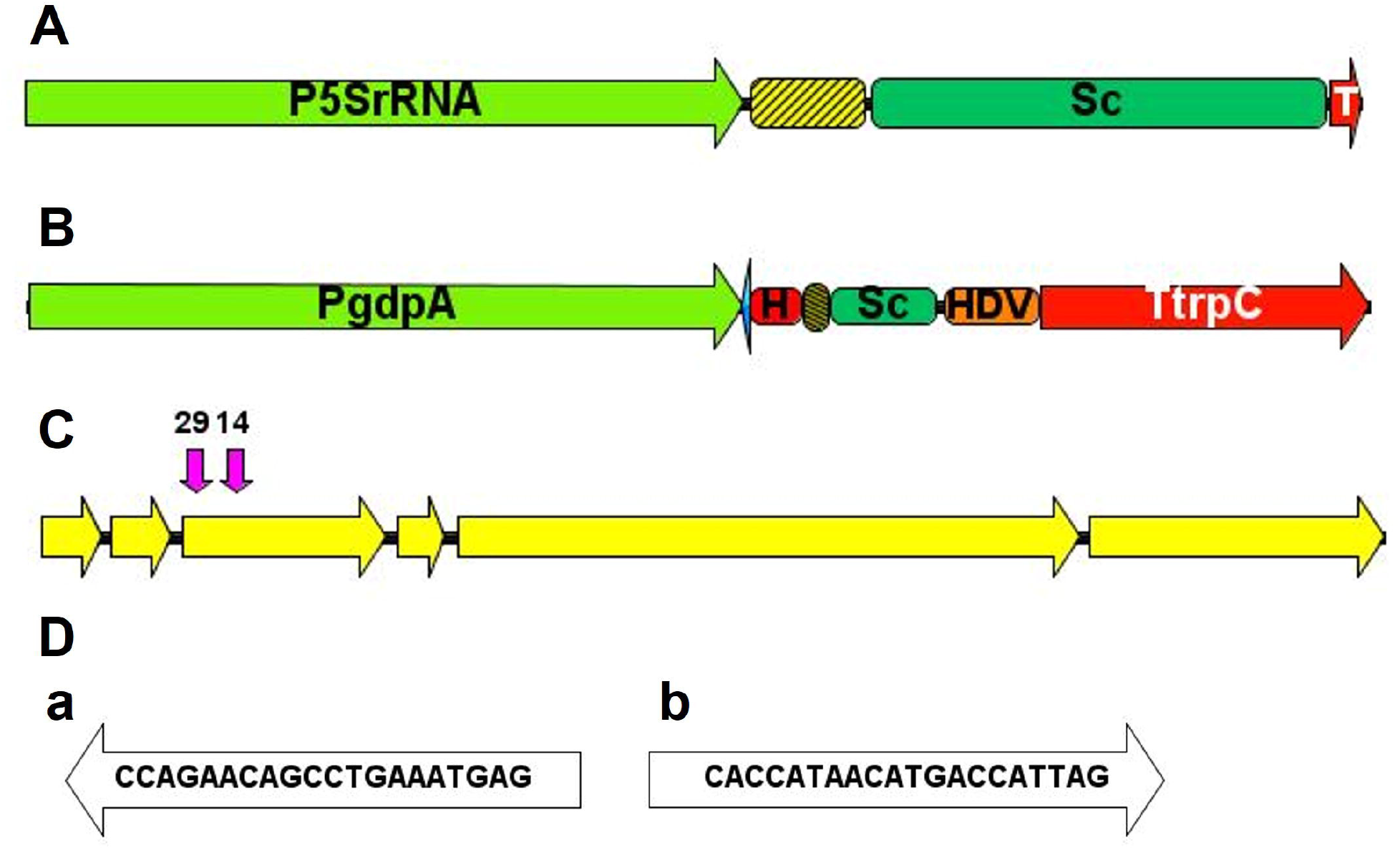
Schematic maps of single guide RNA (sgRNA) expression constructs and the *Fusarium venenatum PKS12* gene highlighting target sites for CRISPR/Cas9 editing. **A** *Fv5SrRNA* promoter-sgRNA construct, showing positions of *PFv5SrRNA*, 20 base target site sequence (yellow and black striped box), scaffold sequence (Sc) and terminator (T). **B** *PgdpA* dual ribozyme-sgRNA construct showing positions of *PgdpA*, inverted repeat (blue arrow) of first six bases in target sequence (yellow and black box), HH and HDV ribozyme sequences and *trpC* terminator (TtrpC). **C** *F. venenatum PKS12* gene showing exons (yellow arrows) and location of the CRISPR target sequences (pink arrows) in exon 3, 3-14 (‘14’) antisense strand, sequence position 3’–5’⍰=⍰238-257 and ‘3-29’ (‘29’) sense strand, position 3’-5’ = 43-62). **D** Target site sequences 3-14 **(a)** and 3-29 **(b)**

### Transformants obtained using CRISPR vectors, phenotypes and sequence analysis of isogenic isolates

AMA1 vectors were maintained in the nucleus under hygromycin selection following PEG-mediated protoplast transformation, and isogenic lines were obtained from colonies developing on hygromycin selection plates by isolation of single spores cultured on non-selective potato sucrose agar (PSA). PSA induces red pigmentation in strains with functional polyketide synthase encoded by the *PKS12* gene and can be used to screen for *PKS12* frameshift variants, which have an albino phenotype (Figure 2). In a preliminary experiment using dual sgRNA vector pFC332::PolII-PK3/14/29, a colony with a clear albino phenotype was obtained indicating disruption of PKS12 protein function. However, using primer pairs which produce an amplicon of 2629 bp from a A3/5 wild type (WT) genomic DNA template, a PCR amplicon of approximately 1 Kb was obtained from variant genomic DNA, for which confusing data was obtained using Sanger sequencing. Results of further PCR analyses, using various primer pairs spanning the target sites, suggest extensive disruption of *PKS12* gene DNA in the variant (Additional file 3: Figure S1).

**Fig. 2.**
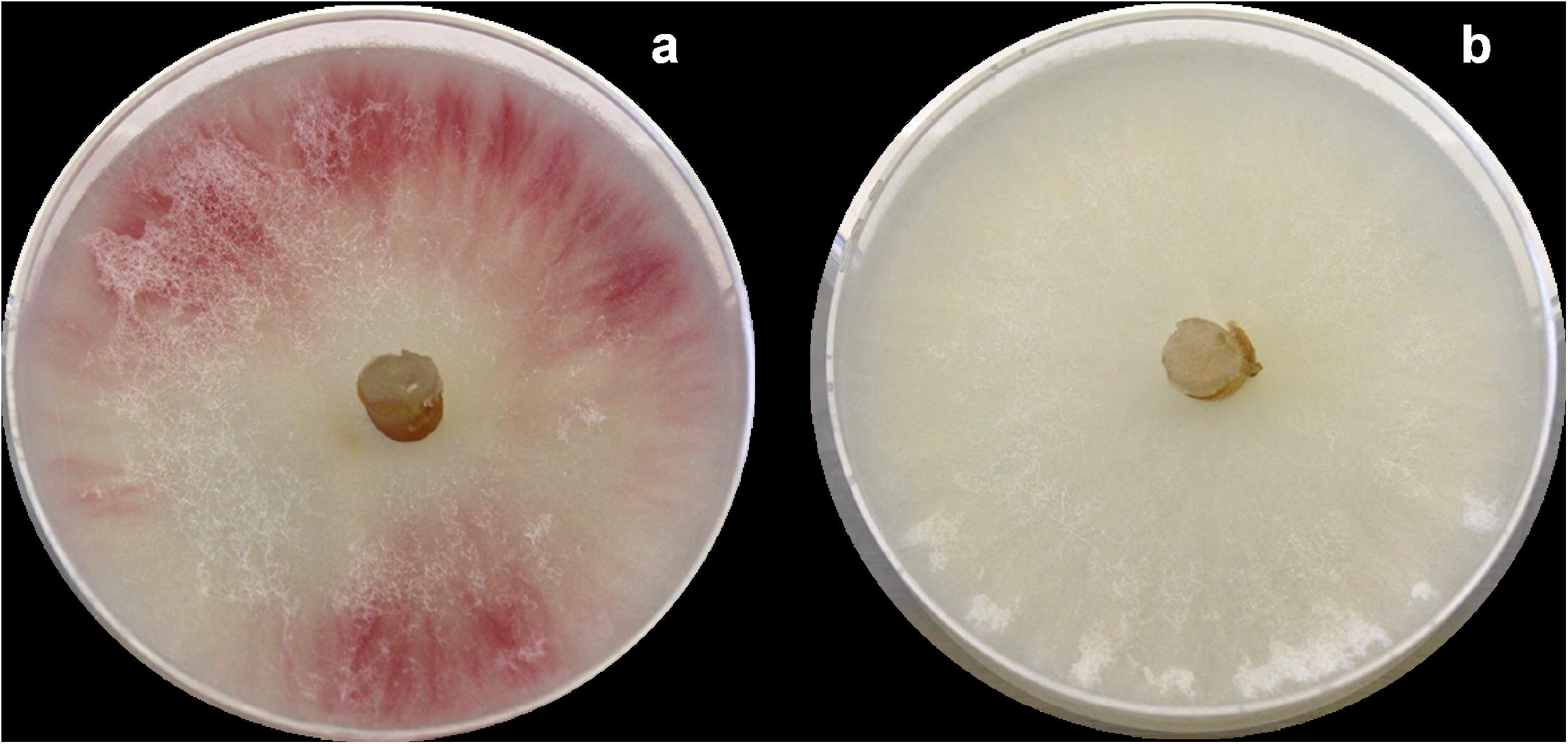
Phenotypes on potato sucrose agar of *F. venenatum* A3/5 **(a)** and *PKS12* gene CRISPR/Cas9 variants (**(b)**

Subsequently a replicated experiment was performed to compare editing efficiency of *Fvcas9* (transcribed from vector pFCFvCas9) with *A. niger cas9* (transcribed from pFC332) and efficacy of sgRNA promoters *PFv5SrRNA* and *PgdpA*, using single sgRNA vectors pFCFvCas9::5S-PK3/14, pFC332::5S-PK3/14 and pFC332::PolII-PK3/14. For each experiment three replicate transformations were performed with each vector and after 10 days the number of mycelial colonies observed on transformation plates was recorded (Table 1). A total of only 2 colonies developed from protoplasts transformed with empty vector pFCFvCas9, compared to 35 from protoplasts transformed with empty vector pFC332 and only one colony developed from protoplasts transformed with pFCFvCas9::PolII-PK3/14. Using vector pFC332, fewer colonies developed when sgRNAs were transcribed from the *Fv5SrRNA* promoter compared to *PgdpA*: a total of 14 transformants were obtained using pFC332::5S-PK3/14 compared to a total of 55 following transformation with pFC332::PolII-PK3/14. Viability of colonies were subsequently assessed (Table 2): viable cultures were not recovered from transformants with vectors pFCFvCas9 or pFCFvCas9::5 SPK-3/14 and viability of transformants co-expressing *A. niger cas9* was less when sgRNAs were transcribed using *PFv5SrRNA* (7 of 12 pFCFvCas9::5SPK-3/14 transformants were viable) compared to transcription from *PgdpA* (all pFC332::PolII-PK3/14 transformants were viable). From each of the viable colonies recovered, at least 10 single spores were cultured on non-selective PSA plates for phenotypic screening. Albino phenotypes were obtained from all of the single spore isolates of pFC332 ::5S-PK3/14 transformed protoplasts, but only 30-40% of isolates of just 3 of the 11 colonies from pFC332::PolII-PK3/14 transformed protoplasts (Table 3). Sequence data from target site amplicons show a single adenine base insertion at position ^-^1 of the cut site in variants (Additional File 4: Table S3), except isolates from colony 5-7 from pFC332::PolII-PK3/14 transformed protoplasts for which no evidence of target site indels was obtained.

**Table 1.**
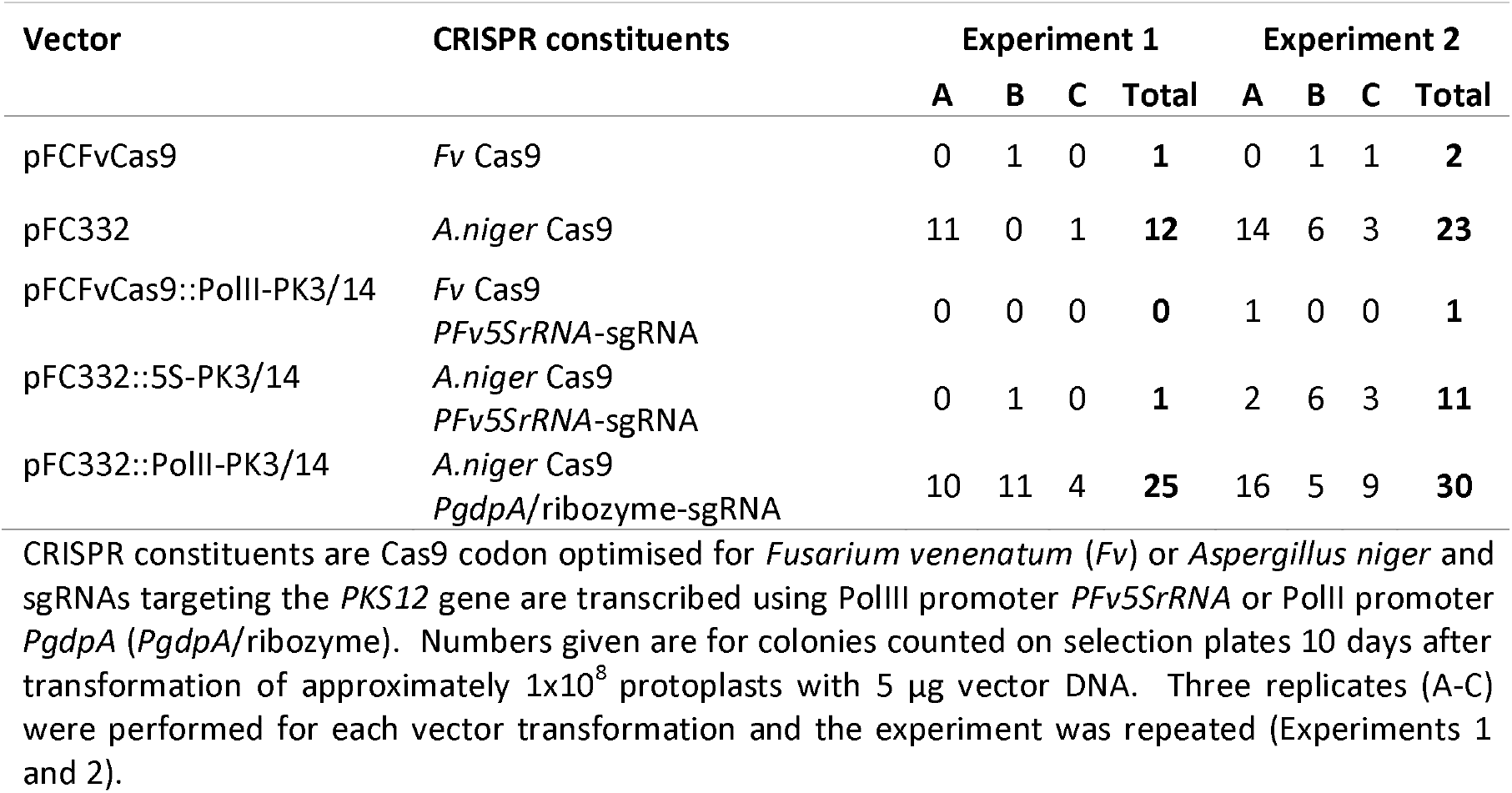
Numbers of colonies regenerating from protoplasts of *F. venenatum* transformed with AMA1 ‘CRISPR’ vectors.

**Table 2.** Growth and viability of *F. venenatum* colonies after transformation with AMA1 CRISPR vectors.

| Vector | Transformed colonies sub-cultured | Growth on selection | Viability on non-selective medium |
| --- | --- | --- | --- |
| pFCFvCas9 | 3 | 0 | 0/3 |
| pFC332 | 3 | 2 | 0/1 |
| pFCFvCas9::PolII-PK3/14 | 1 | 0 | 0/1 |
| pFC332::5S-PK3/14 | 12 | 7 | 0/5 |
| pFC332::PolII-PK3/14 | 27 | 25 | 2/2 |

Viability of colonies from protoplasts transformed with empty CRISPR vectors pFCFvCas9 and pFC332 (incorporating constitutive expression cassettes for *cas9* codon optimised for *F.*
*venenatum* (Fv) and *A. niger*), and vectors with single sgRNA constructs transcribed from the *Fv5SrRNA* promoter (pFC332::5S-PK3/14) or the *PgdpA*-dual ribozyme cassette (pFCFvCas9::PolII-PK3/14 and pFC332::PolII-PK3/14). Colonies from transformation plates were sub-cultured on to selective plates and scored for growth. After 14 days those showing no growth were transferred to non-selective plates and assessed further for viability.

**Table 3.** Phenotypes on PSA of isolates from putative *PKS12* gene variants.

| Putative variant | Number of isolates | Phenotype |  | % albino phenotypes |
| --- | --- | --- | --- | --- |
|  |  | Red | Albino |  |
| pFC332 control | 4 | 4 | - | 0 |
| 5S-1 | 7 | - | 7 | 100 |
| 5S-2 | 7 | - | 7 | 100 |
| 5S-4 | 10 | - | 10 | 100 |
| 5S-5 | 9 | - | 9 | 100 |
| 5S-6 | 10 | - | 10 | 100 |
| 5S-7 | 10 | - | 10 | 100 |
| 5S-10 | 10 | - | 10 | 100 |
| PolII-1 | 12 | 12 | - | 0 |
| PolII-2 | 9 | 9 | - | 0 |
| PolII-3 | 9 | 9 | - | 0 |
| PolII-4 | 14 | 14 | - | 0 |
| PolII-5 | 10 | 7 | 3 | 30 |
| PolII-6 | 10 | 6 | 4 | 40 |
| PolII-7 | 11 | 7 | 4 | 36 |
| PolII-8 | 10 | 10 | - | 0 |
| PolII-9 | 10 | 9 | - | 0 |
| PolII-12 | 10 | 9 | - | 0 |
| PolII-13 | 10 | 10 | - | 0 |

Putative *PKS12* gene variants were generated using sgRNAs transcribed from the PolIII promoter *PFv5SrRNA* (5S) or the PolII promoter *PgdpA* (PolII). Control vector pFC332 is without sgRNA constructs. Isogenic isolates were cultured on potato sucrose agar (PSA), which induces red pigmentation in strains with functional polyketide synthase encoded by the *PKS12* gene. *PKS12* frameshift variants have an albino phenotype.

### AMA1 vector persistence in absence of selection

To test for AMA1 vector persistence in the absence of hygromycin selection, plates with PDA and selective PDA (supplemented with hygromycin B) were inoculated with agar plugs from isogenic *PKS12* variants, which had been maintained for two passages on non-selective media. For comparison, the same media were also inoculated with single spore isolates of mycelium transformed with recombinant AMA1 vectors expressing *mEGFP* instead of CRISPR constituents, which had been maintained on hygromycin selection (Table 4). As expected, control A3/5 wild type (WT) mycelium grew on PDA but not selective PDA and mycelium with AMA1 *mEGFP* recombinant vectors were able to grow on selective medium. All *PKS12* variant isolates grew on PDA but not selective medium, except isolates from three transformants generated using pFC332::PolII-PK3/14: two of three isolates from one transformant and all three isolates from two of the transformants were able to grow on selection, indicating *hph* gene expression from vector DNA.

**Table 4.** Persistence of hygromycin tolerance in CRISPR variants.

| CRISPR variant | Growth on PDA | Growth on PDA+Hyg |
| --- | --- | --- |
| 5S-1 | 3 | 0 |
| 5S-2 | 3 | 0 |
| 5S-4 | 3 | 0 |
| 5S-5 | 3 | 0 |
| 5S-6 | 3 | 3 |
| 5S-7 | 3 | 2 |
| 5S-10 | 3 | 3 |
| PolII-5 | 3 | 0 |
| PolII-6 | 3 | 0 |
| PolII-7 | 3 | 0 |
| Control cultures |  |  |
| AMA1-A | y | y |
| AMA1-B | y | y |
| WT | y | x |

Viability (growth) on hygromycin (selective) PDA (PDA+Hyg) of isogenic lines from CRISPR *PKS12* gene variants using inoculum from cultures previously maintained on non-selective media for two culture passages. Results are shown for 3 isogenic lines from each variant. Variants were generated using sgRNAs transcribed from the PolIII promoter *PFv5SrRNA* (5S) or the PolII promoter *PgdpA* (PolII) respectively. Control cultures transformed with AMA1 vectors expressing *mEGFP* (AMA1-A and AMA1-B) were maintained on selection medium for several culture passages and growth for three replicates of each is denoted by y (growth) or x (no growth).

## Discussion

This is the first report of CRISPR/Cas mediated gene editing in the industrially important fungus *F. venenatum*, used in production of mycoprotein. We demonstrate efficient transgene-free target site mutation of the endogenous marker gene *PKS12* by transient expression of CRISPR constituents from an AMA1 replicator vector.

Use of an autonomously replicating vector for transient expression of CRISPR reagents and selection markers avoids the difficulties associated with *in-vitro* generated unstable sgRNAs, making this an attractive system for generating transgene-free improved strains of *F. venenatum*. Our results indicate loss of the vector in the absence of selection in most isolates, demonstrated by an inability to grow on hygromycin selection media. The continuing hygromycin resistance observed in a small number of isolates suggests either possible chromosomal integration of vector elements (including the *hph* gene for hygromycin resistance), or residual extrachromosomal vector (which may be due to higher initial copy number within some isolates). The recovery of both hygromycin resistant and susceptible single spore isolates derived from one transformation colony suggests loss of residual vector from some but not all nuclei prior to spore formation. Eventual loss from all nuclei would be expected through continued culture in non-selective conditions, as reported by Leeuwe *et al* (19) or after repeated single spore isolation (5).

High efficiency editing is a prerequisite for enabling low input screening of transformants to identify target site variants and may be achieved by optimising levels of sgRNA transcription and nuclear-localised Cas9 codon optimised for the target organism. The correlation between sgRNA transcription levels and efficiency of Cas9-mediated editing was demonstrated by Schwartz in yeast (20), and efficiencies of editing in filamentous fungi have been shown to vary depending on the promoter driving sgRNA transcription and also between different species and experimental systems. Using an AMA1 vector transcribing *A. niger cas9* and single sgRNAs driven by the endogenous PolII promoter *gdpA* within dual ribozymes, Nodvig *et al* (5) reported high efficiency of editing in *A. niger* and *A. brasiliensis*. They were also successful in three other species of *Aspergillus* using the same sgRNA but to varying degrees, which they speculate may be due to various factors including different requirements for *cas9* codon usage and varying levels of *cas9* and sgRNA expression in different species. In a subsequent report Nodvig *et al* (18) noted differences in gene targeting efficiencies between two AMA1 vector selection systems which could be attributed to vector copy number and therefore levels of *cas9* and sgRNA transcription. They also reported higher efficiencies using the *A. fumigatus* PolIII *U3* promoter compared to the polymerase II/ribozyme-based system and suggest this is due to higher sgRNA levels. The effective use of a PolIII promoter driving sgRNA expression was also reported by e.g. Zheng *et al* (17) who achieved high (100%) editing in *Aspergillus* by transcribing sgRNAs from an expression vector using the native *5SrRNA* promoter. Likewise Shi *et al* (14) reported enhanced gene editing in *F. fujikuroi* using promoters for highly expressed endogenous genes: using an expression vector they obtained efficiencies of 79.2% and 37.5% when sgRNAs were transcribed from the *F. fujikuroi* PolIII *5SrRNA* or *U6* promoters, respectively, but were unsuccessful using the heterologous PolII (*PgdpA*) dual ribozyme system. We demonstrate similar results in *F. venenatum* using an AMA1 vector for *in vivo* production of CRISPR reagents, achieving edits in 100% of recovered isolates when sgRNAs were transcribed from the native *Fv5SrRNA* promoter and greatly reduced efficacy using the *PgdpA* ribozyme system including evidence of off-target editing. RT-qPCR results from our preliminary experiments (Additional file 5: Figure S2, Table S4) suggest relatively low levels of transcription from *PgdpA* in AMA1 vector transformants, observed also by Nodvig *et al* (18) in *Aspergillus* expressing *mRFP* from expression vectors.

The importance of the NLS in determining efficiency of Cas9 protein nuclear import and subsequent gene editing success has also been highlighted by several studies. SV40_NLS_ has been used widely for Cas9 nuclear import in eukaryotes including other filamentous fungi (5,6,10,18). Wang *et al* (12) reported that when delivered as a constituent of a RNP, Cas9-SV40_NLS_ was ineffective in *F. oxysporum*, but use of the endogenous NLS from histone H2B was successful. Likewise, Shi *et al* (14) using Cas9 produced *in vitro* compared three NLS in *F. fujikuroi*: SV40_NLS_, the endogenous NLS from histone H2B (HBT_NLS_) which is conserved in eukaryotes and the endogenous NLS from Velvet (VEL_NLS_), a nuclear protein found in many filamentous fungi. They obtained no gene edits using SV40_NLS_ and greatest efficiencies using HBT_NLS_. In our experiments we were able to obtain gene edited variants when *cas9-*SV40_NLS_ was transcribed *in vivo* from the *tef1* promoter using an AMA1 replicator vector and RT-qPCR results show relatively high levels of transcription of *cas9* (Additional file 5: Figure S2, Table S4), indicating that effective gene editing in *Fusarium* mediated by Cas9-SV40_NLS_ is dependent on methodology and sufficient levels of *cas9* transcription.

Cas9 optimization is another key factor reported to influence gene editing success in filamentous fungi, and differences in reported effectiveness and toxicity may be ascribed to codon usage as well as potential differences in levels of nuclear Cas9 resulting from the different methodologies employed. Foster *et al* (10) reported toxicity of Cas9 optimized for both *N. crassa* and the target organism in *M. oryzae* when *cas9* was expressed constitutively from integrated transgenes but were able to recover gene edited mutants using hSpCas9 delivered transiently as RNPs. Fuller *et al* (4) found no evidence of toxicity from hSpCas9 transcribed from the *tef1* promoter in *Aspergillus* transgenic lines, and likewise Shi *et al* (14) observed no effect on growth of *F. fujikuroi* of optimised *FuCas9* transcribed from *PgdpA* using expression vectors. Also, in *F. graminearum* Gardiner and Kazan (11) successfully used *F. graminearum* optimised Cas9 transcribed using the *PgdpA* promoter from integrated transgenes. However, in our experiments using an AMA1 vector for transcription of *cas9* using the *tef1* promoter we were unable to recover viable colonies from protoplasts expressing Cas9 optimised for *F. venenatum* and recovery rates were low after transformation with vectors expressing *A. niger* Cas9, which is indicative of Cas9 toxicity (for example see Additional File 6: Table S5). Additionally, our results show observed reduced viability of protoplasts and mycelium expressing *PFv5SrRNA*-sgRNAs compared to those expressing *PgdpA*-sgRNAs, suggesting a possible interaction between sgRNA transcription levels and Cas9 toxicity.

## Conclusions

In conclusion, we have demonstrated that 100% gene editing frequency in isolates of *F. venenatum* can be achieved by transforming protoplasts with AMA1 replicator vectors expressing *A. niger* codon-optimised *cas9-*SV40_NLS_ transcribed from the *tef1* promoter and sgRNAs transcribed from the endogenous *Fv5SrRNA* promoter. Evidence of AMA1 vector loss in the absence of selection makes this an attractive system for generating transgene-free gene edited strains of *F. venenatum* and for performing successive gene edits using the same selection marker. However, reduced protoplast viability due to potential Cas9 toxicity was observed, which may impede development of further CRISPR/Cas technologies such as gene insertion, base editing and prime editing in *F. venenatum*. Assessment of the efficacy of alternative promoters and/or use of an inducible system for regulating *cas9* expression would enable optimization of Cas9 production and allow re-evaluation of Cas9 codon-optimized for *F. venenatum*, which was apparently toxic when expressed from the *tef1* promoter using an AMA1 vector. Implementation of the gene editing methodology described here, and development of additional CRISPR/Cas tools will further the understanding of controls governing growth and metabolite synthesis in *F. venenatum*. This will enable identification of targets for strain improvement using classical means and facilitate innovation in fermentation development.

## Methods

### Vector construction

Vector backbones were obtained by restriction digest and component parts for vector inserts were generated by PCR from synthesised DNA or existing vector templates using Q5^®^ High-Fidelity 2X Master Mix (NEB). Vector components were purified using the Monarch^®^ DNA Gel Extraction Kit or the Monarch^®^ PCR & DNA Cleanup Kit (NEB) and assemblies were performed using NEBuilder^®^ HiFi DNA Assembly Master Mix (NEB). For details of primers used in vector construction see Additional File 7: Table S6. Recombinant vectors were recovered by transformation of ^®^ NEB 10-beta Competent E. coli (High Efficiency) cells and subsequent plasmid isolation was performed using the NucleoSpin Plasmid kit (Macherey-Nagel).

Codon usage preference of predicted highly expressed genes for *F. venenatum* A3/5 was determined (Additional File 8: Table S7) and used to generate a codon-optimised sequence for Cas9. Using Optimizer (21) ‘Relative Synonymous Codon Usage’ (RSCU) was performed, selecting options for ‘standard genetic code’ and ‘one amino acid - one codon’. The coding sequence was subsequently domesticated to enable use in Golden Gate cloning if required and DNA for the resulting sequence, (Additional File 1: Table S1), was synthesised (GeneArt, Invitrogen). An AMA1 replication vector backbone (pFCBB) of 10,129 bases was obtained from pFC332 (5) by restriction digest using PacI and BamHI-HF^®^ (NEB), excising part of the Nt.BbvCI –PacI cloning site, the *tef1* promoter, *cas9* coding sequence and 489 bases (5’-3’) of the *tef1* terminator. The residual *tef1* terminator (87 bases) was incorporated in recombinant vector assembly by overlap with inserts terminated with *Ttef1*. The SV40_NLS_-stop codon, *tef1* terminator and *tef1* promoter parts were PCR-amplified from template pFC332. *FvCas9* was PCR amplified in one part from synthesised DNA and assembled with the *tef1* promoter and *tef1* terminator parts in Golden Gate vector pICH47751 (22) to give Ptef1-FvCas9-SV40-Ttef1. This was used as a template for generation by PCR of two parts for assembly with pFCBB to give pFCFvCas9: *Ptef1* and *cas9* bases 1-2141 (5’-3’) were amplified as Part 1 using primers NtPac1_Ptef1 (pFCBB)_F and FvCas9-1_R; and Part 2 was generated by primers FvCas9-2_F and Ttef1_R_no ext, incorporating *cas9* bases 2142-4376 (5’-3’) and sequences for SV40-stop codon and *Ttef1*. The Nt.BbvCI –PacI cloning site was reconstructed by incorporating required bases into the 5’ extension of primer NtPac1_Ptef1 (pFCBB)_F. The PolII/ribozyme sgRNA cassettes were constructed using pFC334 as a template and assembled into the Nt.BbvCI-PacI USER site of pFC332 as described by Nodvig *et al* (5). *5SrRNA* sgRNA cassettes were synthesised by IDT (Additional File 2: Table S2) and PCR amplified using primers 5SrRNA_F USER and 5SrRNA_R USER for USER assembly into pFC332 or pFCBB. Details of USER reactions are described in Additional File 9: Table S10.

### Protoplast transformation

Protoplast transformation was performed as described in Additional File 10: Table S11. Media used are detailed in Additional File 11: Table S12. Colonies from transformation plates were transferred after 10 days to PDA supplemented with hygromycin B 75 µg/ml, then cultured for 2 weeks at 28 °C. For single spore isolation agar plugs were inoculated in 20 ml CMC medium in a 100 ml Erlenmeyer flask and incubated shaking at 175 rpm for 7 days at 25 °C. Spores were collected by filtration through 2 layers of Miracloth (Millipore), pelleted at 4000xg for 10 minutes and plated to give a low density on PDA plates. After incubation for 14-15h at 20 °C single germinating spores were isolated and incubated in the dark at 28 °C on PSA (Additional file 12: Table S13) for identifying albino *PKS12* variants.

### CRISPR variant analysis

Genomic DNA for use in PCR was extracted from mycelium cultured in 50 ml plastic conical based tubes containing 10 ml of a proprietary medium and incubated shaking at 175 rpm for 5-7 days at 28 °C. The culture was collected on Miracloth, rinsed with autoclaved purified water, blotted to remove excess liquid and added with 2 ball bearings (4 mm chrome steel) to 2 ml microcentrifuge tubes before flash freezing in liquid nitrogen and storage at ^-^80 °C. Mycelium was ground while frozen using a ball mill (Retsch) and DNA was extracted using the DNeasy Plant Mini Kit (Qiagen). PCR amplicons were generated using Q5^®^ High-Fidelity 2X Master Mix (NEB) using primers PKS12_F2 and PKS12_R2, which span the target region (Figure 1). Amplicons were Sanger sequenced (Eurofins Genomics) using sequencing primers PKS12-Seq_R1 and PKS12-Seq_F2.

### Sequencing analysis and software

Nucleotide and protein alignments were performed using Geneious version 10.0.2 (http://www.geneious.com) (23). Schematic maps were prepared using IBS software (24).

## Abbreviations

Cas9: CRISPR associated DNA-cutting enzyme
CRISPR: clustered regularly interspaced short palindromic repeats
HDV ribozyme: Hepatitis Delta Virus ribozyme
HH ribozyme: Hammerhead ribozyme
indel: insertion-deletion mutation of genomic sequence
Kb: Kilobase pairs
mEGFP: monomeric enhanced green fluorescent protein
mRFP: monomeric red fluorescent protein
PDA: Potato Dextrose Agar
PolII: polymerase II
PolIII: polymerase III
USER: Uracil-Specific Excision Reagent
PCR: polymerase chain reaction
RT-qPCR: reverse transcription quantitative PCR

## Additional Files

**Additional File 1: Table S1**. Expression cassette Ptef1-FvCas9-SV40-Ttef1 used in this study.

**Additional File 2: Table S2**. Sequence of *PFv5SrRNA*-sgRNA cassettes used in this study.

**Additional File 3: Figure S1**. Schematic map of 5’ exons of the *F. venenatum PKS12* gene, showing CRISPR/Cas9 target sites and primers used in analysis of dual-sgRNA variant genomic DNA.

**Additional File 4: Table S3**. *PKS12* gene target site sequence data for phenotypic variants.

**Additional File 5: Figure S2**. RT-qPCR analysis of *F. venenatum* AMA1 vector *cas9* transcripts expressed in *F. venenatum*; **Table S4**. Primers used in RT-qPCR analysis.

**Additional File 6: Table S5** Viability of protoplasts transformed with AMA1 vectors expressing *mEGFP* and *cas9*

**Additional File 7: Table S6**. Primers used in vector construction and PCR analysis of PKS12 gene variants.

**Additional file 8: Table S7**. Codon usage table for highly expressed genes in *F. venenatum*.

**Additional File 9: Table S10**. Method for USER cloning of promoter-sgRNA cassettes into AMA1 vectors.

**Additional File 10: Table S11**. Method used for protoplast isolation and transformation.

**Additional File 11: Table S12**. Media used for making protoplasts and protoplast transformation.

**Additional File 12: Table S13**. Method for making Potato Sucrose Agar (PSA).

## Ethics approval and consent to participate

Not applicable.

## Consent for publication

Not applicable.

## Competing interests

The authors declare that they have no competing interests.

## Funding

This work was supported by the Biotechnology and Biological Sciences Research Council (BBSRC) and (BB/P020364/1) and Marlow Foods.

## Authors’ contributions

FMW and RJH devised the research plan. FMW planned the experiments, designed and constructed the CRISPR cassettes and vectors, performed transformation experiments and conducted phenotypic and molecular analyses of variants. FMW wrote the manuscript with input from RJH and Marlow Foods. Both authors reviewed the manuscript. Both authors read and approved the final manuscript.

## Acknowledgments

The authors thank the following: Marlow Foods for financial support and participation at project inception and throughout the duration of the project, and for provision of strains and technical advice; the vectors pFC332 (Addgene Plasmid #87845) and pFC334 (Addgene Plasmid #87846) were a gift from the Uffe Mortensen laboratory, Technical University of Denmark; synthesised DNA for *Fvcas9* was kindly provided through the services of SCLU; A.D. Armitage annotated the *F. venenatum* genome and produced the codon usage table for *F. venenatum*; J Bennett performed RNA extractions and qRT-PCRs including data analysis.

## Notes

### Competing Interest Statement

The authors have declared no competing interest.

