## Additional File 3 for "CRISPR/Cas9 mediated editing of the Quorn fungus *Fusarium venenatum* A3/5 by transient expression of Cas9 and sgRNAs targeting endogenous marker gene *PKS12*"

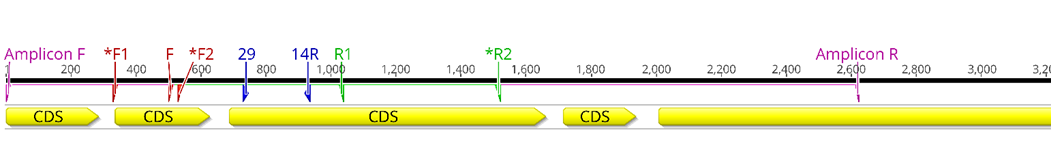

**Fig. S1**	Schematic map of 5’ exons of the *Fusarium venenatum* *PKS12* gene, showing CRISPR/Cas9 target sites and primers used in analysis of dual-sgRNA variant genomic DNA. Yellow arrows are exons (CDS). Blue arrows are target sites ‘29’ (left) and ’14’ (right) in the third exon. Pink primers (‘Amplicon F’ and ‘Amplicon R’) were used to amplify a product for sequence analysis of target site variants: amplicons generated were 2629 bp from WT genomic DNA and approximately 1Kb from variant DNA. Sanger sequencing of amplicons gave expected sequence for the WT amplicon using primers ‘*F1’, ‘*F2’ and ‘*R2’. However, for the variant amplicon identifiable sequence data were obtained only for the sense strand of the first exon using primers ‘F1’, ‘F2’ and ‘R2’. Expected PCR products were generated from WT DNA using primers ‘F’ and ‘R1’ or ‘R2’, but it was not possible to generate PCR products from variant DNA using ‘R1’ or ‘R2’ with any of forward primers ‘Amplicon F’, ‘F1’ or ‘F2’.
