## Additional File 4 for "CRISPR/Cas9 mediated editing of the Quorn fungus *Fusarium venenatum* A3/5 by transient expression of Cas9 and sgRNAs targeting endogenous marker gene *PKS12*"

**Table S3 *PKS12* gene target site sequence data for phenotypic variants**

| Reference amino acid sequence | | | PLISGCSGEEFQPLSYTDLLHCCVTDM |
| --- | --- | --- | --- |
| Variant amino acid sequence | | | PLNFRLFWGRIPAPELYRPPPLLCD*H |
| Reference DNA sequence | | | **CCCCTCATTTCAGGCTGTTCTGG**GGAAGAATTCCAGCCCCTGAGCTATACCGACCTCCTCCACTGCTGTGTGACTGACATG |
| DNA sequences for phenotypic variants | | | |
| 5S | 1 | a | CCCCTC**A**ATTTCAGGCTGTTCTGGGGAAGAATTCCAGCCCCTGAGCTATACCGACCTCCTCCACTGCTGTGTGACTGACAT |
| 5S | 1 | b | CCCCTC**A**ATT***c***TCAGGCTGTTCTGGGGAAGAATTCCAGCCCCTGAGCTATACCGACCT***a***CCTaCCACT***c***GCTG***ggg***GTGACT |
| 5S | 1 | c | CCCCTC**A**ATTTCAGGCTGTTCTGGGGAAGAATTCCAGCCCCTGAGCTATACCGACCTCCTCCACTGCTGTGTGACTGACAT |
| 5S | 2 | a | Failed |
| 5S | 2 | b | CCCCTC**A**ATTTCAGGCTGTTCTGGGGAAGAATTCCAGCCCCTGAGCTATACCGACCTCCTCCACTGCTGTGTGACTGACAT |
| 5S | 2 | c | CCCCTC**A**ATTTCAGGCTGTTCTGGGGAAGAATTCCAGCCCCTGAGCTATACCGACCTCCTCCACTGCTGTGTGACTGACAT |
| 5S | 4 | a | CCCCTC**A**ATTTCAGGCTGTTCTGGGGAAGAATTCCAGCCCCTGAGCTATACCGACCTCCTCCACTGCTGTGTGACTGACAT |
| 5S | 4 | b | CCCCTC**A**ATTTCAGGCTGTTCTGGGGAAGAATTCCAGCCCCTGAGCTATACCGACCTCCTCCACTGCTGTGTGACTGACAT |
| 5S | 4 | c | CCCCTC**A**ATTTCAGGCTGTTCTGGGGAAGAATTCCAGCCCCTGAGCTATACCGACCTCCTCCACTGCTGTGTGACTGACAT |
| 5S | 5 | a | CCCCTC**A**ATTTCAGGCTGTTCTGGGGAAGAATTCCAGCCCCTGAGCTATACCGACCTCCTCCACcGCTGTGTGACTGACAT |
| 5S | 5 | b | CCCCTC**A**ATTTCAGGCTGTTCTGGGGAAGAATTCCAGCCCCTGAGCTATACCGACCTCCTCCACTGCTGTGTGACTGACAT |
| 5S | 5 | c | CCCCTC**A**ATTT***t***CAGGCTGTTCTGGGGAAGAATTCCAGCCCCTGAGCTATACCGACCTCCTCCAC***c***GCTGTGTGAC***c***GACA***c*** |
| 5S | 6 | a | CCCCTC**A**ATTTCAGGCTGTTCTGGGGAAGAATTCCAGCCCCTGAGCTATACCGACCTCCTCCACTGCTGTGTGACTGACAT |
| 5S | 6 | b | Failed |
| 5S | 6 | c | CCCCTC**A**ATTTCAGGCTGTTCTGGGGAAGAATTCCAGCCCCTGAGCTATACCGACCTCCTCCAC***c***GCTGTGTGACTGACAT |
| 5S | 7 | a | Failed |
| 5S | 7 | b | CCCCTC**A**ATTTCAGGCTGTTCTGGGGAAGAATTCCAGCCCCTGAGCTATACCGACCTCCTCCACTGCTGTGTGACTGACAT |
| 5S | 7 | c | Failed |
| 5S | 10 | a | Failed |
| 5S | 10 | b | CCCCTC**A**ATTTCAGGCTGTTCTGGGGAAGAATTCCAGCCCCTGAGCTATACCGACCTCCTCCACaGCTGTGTGACTGACAT |
| 5S | 10 | c | CCCCTC**A**ATTTCAGGCTGTTCTGGGGAAGAATTCCAGCCCCTGAGCTATACCGACCTCCTCCACTGCTGTGTGACTGACAT |
| PolII | 5 | a | CCCCTC**A**ATTTCAGGCTGTTCTGGGGAAGAATTCCAGCCCCTGAGCTATACCGACCTCCTCCACTGCTGTGTGACTGACAT |
| PolII | 5 | b | Failed |
| PolII | 5 | c | CCCCT***t***C**A**ATTTCAGGCTGTTCTGGGGAAGAATTCCAGCCCCTGAGCTATACCGACCTCCTCCAC***a***GCTGTGTGACTGACAT |
| PolII | 6 | a | CCCCTC**A**ATTTCAGGCTGTTCTGGGGAAGAATTCCAGCCCCTGAGCTATACCGACCTCCTCCAC***a***GCTGGGTGACTGACAT |
| PolII | 6 | b | CCCCTC**A**ATTTCAGGCTGTTCTGGGGAAGAATTCCAGCCCCTGAGCTATACCGACCTCCTCCACTGCTGTGTGACTGACAT |
| PolII | 6 | c | CCCCTC**A**ATTTCAGGCTGTTCTGGGGAAGAATTCCAGCCCCTGAGCTATACCGACCTCCTCCAC***a***GCTG***g***GTGACTGACAT |
| PolII | 6 | d | CCCCTC**A**ATTTCAGGCTGTTCTGGGGAAGAATTCCAGCCCCTGAGCTATACCGACCTCCTCCACTGCTGTGTGACTGACAT |
| PolII | 7 | a | CCCCTCATTTCAGGCTGTTCTGGGGAAGAATTCCAGCCCCTGAGCTATACCGACCTCCTCCAC***c***GCTGTGTGACTGACAcG |
| PolII | 7 | b | CCCCTCATTTCAGGCTGTTCTGGGGAAGAATTCCAGCCCCTGAGCTATACCGACCTCCTCCACTGCTGTGTGACTGACATG |
| PolII | 7 | c | CCCCTCATTTCAGGCTGTTCTGGGGAAGAATTCCAGCCCCTGAGCTATACCGACCTCCTCCACTGCTGTGTGACTGACATG |
| PolII | 7 | d | CCCCTCATTTCAGGCTGTTCTGGGGAAGAATTCCAGCCCCTGAGCTATACCGACCTCCTCCACTGCTGTGTGACTGACATG |

*PKS12* gene variants were generated using sgRNAs transcribed from the PolIII promoter *PFv5SrRNA* (5S) or the PolII promoter *PgdpA* (PolII). Samples shown are numbered to indicate separate colonies recovered from selection plates following protoplast transformation. Lower font letters identify separate isogenic lines (all with an albino phenotype) derived from colonies. Sequence data (5’-3’) for the sense strand are shown for 90 bases. Reference DNA sequence corresponds to data obtained from genomic DNA of the wild type and colonies recovered from protoplasts transformed with ‘empty vector’ pFC332 (red font corresponds to the target site on the antisense strand). The bases corresponding to the PAM sequence (GGG) on the antisense strand are underlined (CCC). In all variants except isolates from colony ‘7’ a single base insertion (‘**A**’) is seen at position -1 of the ‘cut site’, which is 3 base pairs upstream of the PAM site on the antisense strand. This causes a frameshift generating variant amino acid sequence and a stop codon (*****) within exon 3, shown at the top of the table. Target site sequence is close to the 5’ end of the reads obtained, in some samples either generating what is taken to be spurious data (lower case font), or resulting in omission of target site sequence from the read (‘Failed’).
