## Additional File 5 for "CRISPR/Cas9 mediated editing of the Quorn fungus *Fusarium venenatum* A3/5 by transient expression of Cas9 and sgRNAs targeting endogenous marker gene *PKS12*"

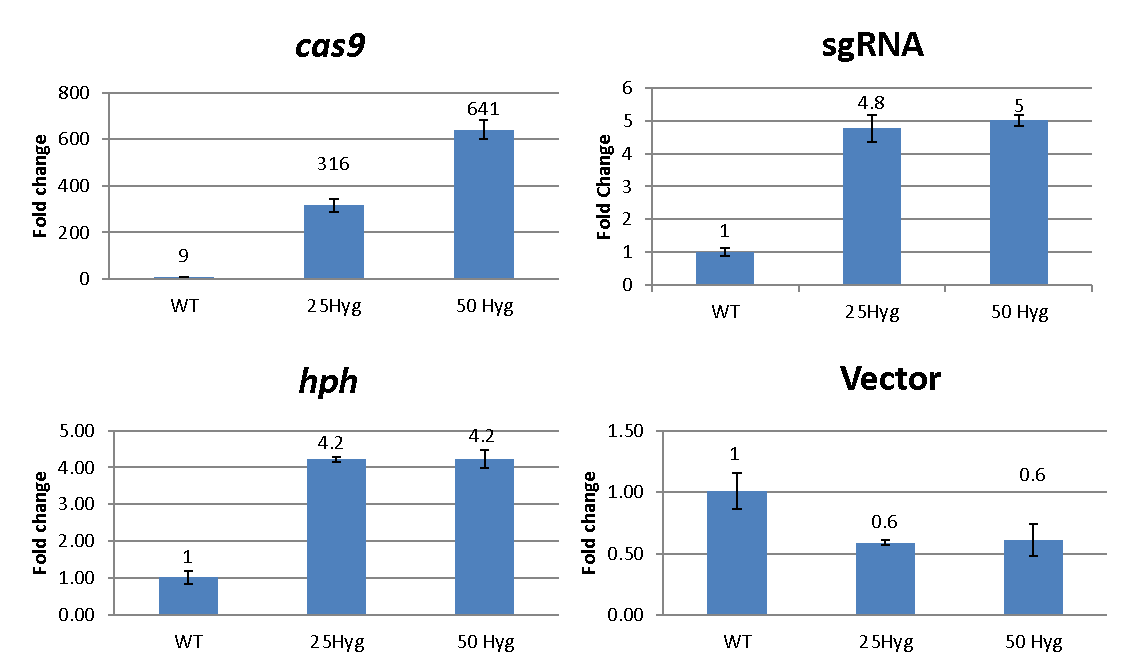

**C**

**D**

**B**

**A**

**Fig. S2** qRT-PCR analysis of *F. venenatum* AMA1 vector *cas9* transcripts in *F. venenatum*. **A** *cas9* transcripts driven from the *tef1* promoter. **B** sgRNA transcript driven from *PgdpA.* **C** Transcription of the *hph* gene for hygromycin selection of vectors. **D** Primers spanning the *Ptef1*-*cas9* gene junction were included to give a baseline for comparison with AMA1 vector transcripts. AMA1 vector was maintained in mycelium using hygromycin at 25 µg/ml (25Hyg) or 50 µg/ml (50Hyg). Wild type mycelium (WT) was cultured in non-selective medium. Mycelium was incubated at 28 °C shaking at 180 rpm, in a proprietary medium for 14 days before collection for total RNA extraction and cDNA synthesis. Primers used for quantification of vector genes and three fungal housekeeping reference genes used for normalisation of data are listed in Table S4. Relative quantification was calculated by the deriving the ratio (ΔCt) between the amounts (the Ct value) of the target gene and the geomean of the reference. Bars represent the means and standard errors of three biological replicates. Mean values are shown above each bar.

**Table S4 Primers used in qRT-PCR analysis of AMA1 vector and *tri5* gene transcripts**

|  | Primer sequence | Amplification product |
| --- | --- | --- |
| *Housekeeping reference genes* |  |  |
| EF1b-2F | CCTCCAGGATGTCTACAAGA | *elongation factor-1* transcript |
| EF1b-2R | CTCAACGGACTTGACTTCAG |  |
| bTUBa-1F | GTTGATCTCCAAGATCCGTG | *β-tubulin* transcript |
| bTUBa-1R | CATGCAAATGTCGTAGAGGG |  |
| GADPH_F | TGACTTGACTGTTCGCCTCGAGAA | *GAPDH* transcript |
| GADPH_R | ATGGAGGAGTTGGTGTTGCCGTTA |  |
| *AMA1 vector* |  |  |
| Ptef1-Cas9_F | TCGCTTCTCTCCTCCATCCT | *Ptef1*-*cas9* gene junction |
| AMA1-Ptef1_R | CCGTAATCACAGCCCAACCA |  |
| qPCR-Cas9_F | GCCCAAATCGGTGACCAGTA | *cas9* transcript |
| qPCR-Cas9_R | CTTCGCGATTCAGCTTGACG |  |
| qPCR_sgRNA_F | CCCCCGAACGTTTTAGAGCT | sgRNA transcript |
| qPCR_sgRNA_R | CGACTCGGTGCCACTTTTTC |  |
| qPCR-Hyg_F | CTCGGAGGGCGAAGAATCTC | *hph* transcript |
| qPCR-Hyg_R | TCGCTGAACTCCCCAATGTC |  |

Housekeeping gene primers are taken from Harris LJ *et al* (2016) doi.org/10.1016/j.funbio.2015.10.010.
