## Additional File 6 for "CRISPR/Cas9 mediated editing of the Quorn fungus *Fusarium venenatum* A3/5 by transient expression of Cas9 and sgRNAs targeting endogenous marker gene *PKS12*"

**Table S5 Viability of protoplasts transformed with AMA1 vectors expressing *mEGFP* and *cas9***

| Protoplast-derived colonies | Selection plate |
| --- | --- |
| A - pFC::PgpdA-mEGFP-SV40-tTrpC | 110 |
| B - pFC332:: PgpdA-mEGFP-SV40-tTrpC (Cas9) | 7 |

Number of colonies observed on hygromycin selection plates following transformation of protoplasts with AMA1 vectors expressing *mEGFP* (A) or *mEGFP* and *A. niger cas9* (B).
