## Additional File 7 for "CRISPR/Cas9 mediated editing of the Quorn fungus *Fusarium venenatum* A3/5 by transient expression of Cas9 and sgRNAs targeting endogenous marker gene *PKS12*"

**Table S6 Primers used in vector construction and PCR analysis of *PKS12* gene variants**

| Primers/*construct*/*vector* | Primer sequence 5’-3’ | Part generated/primer function |
| --- | --- | --- |
| *pICH47751::Ptef1-FvCas9-SV40-stop-Ttef1* * | | |
| pTef1 (47751)_F | aaactagaattcgagctcCGAGACAGCAGAATCACCGC | Ptef1 |
| pTef1 (FvCas9)_R | cttcttgtccattGGTGAAGGTTGTGTTATGTTTTGTG |  |
| FvCas9 (pTef1)_F | accttcaccATGGACAAGAAGTACTCTATCGGC | FvCas9 |
| FvCas9 (tTef1)_R | cttgggaggGTCGCCGCCGAGCTG |  |
| SV40-STOP-tTef1 (FvCas9)_F | ggcggcgacCCTCCCAAGAAGAAGCGCAA | SV40-stop-Ttef1 |
| tTef1 (47751)_R | cgtgcagaagacaagtaaGTATTGGGATGAATTTTGTATGCACG |  |
| *pFCFvCas9* * | | |
| NtPac1_Ptef1 (pFCBB)_F | ttccgctgagggtttaattaagacctcagcCGAGACAGCAGAATCACCGC | Ptef1-FvCas9 5' |
| FvCas9-1_R | tcgccctggCCAGAAACCTGAGCCTTCTGG |  |
| FvCas9-2_F | ggtttctggCCAGGGCGACTCTCTCCA | FvCas93'-SV40-stop-Ttef1 |
| tTef1R(pFCBB)long | GCTCACCGCCTGGACGACTAAACCAAAATAGGCATTGATGTGTTGACCTCCACTAGCATTACACTTGTATTGGGATGAATTTTGTATGCACG |  |
| *PollII/ribozyme sgRNA constructs USER primers* ** | | |
| PKS3-14R_F | *AGTAAGCUCGT*CCCAGAACAGCCTGAAATGAGGTTTTAGAGCTAGAAATAGCAAGTTAAA | PolII-PK3/14 |
| PKS3-14R_R | *AGCTTACUCGTTTCGTCCTCACGGACTCATCAG*CCAGAACGGTGATGTCTGCTCAAGCG |  |
| PKS3-29_F | *AGTAAGCUCGT*CCACCATAACATGACCATTAGGTTTTAGAGCTAGAAATAGCAAGTTAAA | PolII-PK3/29 |
| PKS3-29_R | *AGCTTACUCGTTTCGTCCTCACGGACTCATCAG*CACCATCGGTGATGTCTGCTCAAGCG |  |
| *P5SrRNA constructs USER primers* *** | | |
| 5SrRNA_F USER | gggtttaauCACATACGACCAAAGGTAGTGGA | 5S-PK3/14 and 5S-PK3/29 |
| CSN390 {Nodvig 2015} | ggtcttaauGAGCCAAGAGCGGATTCCTC |  |
| 5SrRNA_R USER | ggtcttaauTGATCCATGCACTCCGGGT |  |
| CSN389 {Nodvig 2015} | gggtttaauGCGTAAGCTCCCTAATTGGC |  |
| *Primers for analysis and sequencing* |  |  |
| FvCas9_int_F | CTGTCGAGATCTCTGGCGTC | Amplicon of *Fvcas9* junction (pFCFvCas9) |
| FvCas9_int_R | TGTCAGACTTGCCTCGGTTC |  |
| FvCas9-Seq_F | ACGACAAGGTCATGAAGC | Sequencing primers for *Fvcas9* internal ligation junction |
| FvCas9-Seq_R | CTGGTCGACGTACATGTC |  |
| PKS12-F-2 | GGGTATGGCGAACTACCTCG | Amplicon spanning CRISPR/Cas9 target sites‘14’ & ‘29’ in PKS12 gene exon 3 |
| PKS12-R-2 | CATAAGCTGTTTCGAGCGCC |  |
| PKS12-Seq_F2 | CTATGCTCCGTACCATGC | Sequencing primers for target site amplicons |
| PKS12-Seq_R1 | TCTTCAAGTCCTGGCGCTG |  |

* Sequences in lower case font are primer extensions for Gibson assembly. ** L to R: italicised font is USER overhang Hammerhead ribozyme sequence (3' forward primers, 5' reverse primers). The uracil base in the USER overhang is italicised and underlined. Bold highlighted font is target sequence; the first 6 bases are underlined. Capital font which is not italicised or underlined is sequence binding to sgRNA scaffold (forward primers) or PgdpA (reverse primers). *** Sequences in lower case font are primer extensions for USER cloning into pFC332 or pFCBB, capital font is sequence for 5SrRNA promoter 5' (forward primer) or spacer at 3' end of the 5SrRNA cassette
