## Additional File 8 for "CRISPR/Cas9 mediated editing of the Quorn fungus *Fusarium venenatum* A3/5 by transient expression of Cas9 and sgRNAs targeting endogenous marker gene *PKS12*"

**Table S7 Codon usage table for highly expressed genes in *F. venenatum***

UUU: 0.76; UCU: 1.28; UAU: 0.70; UGU: 0.78; UUC: 1.24; UCC: 1.08; UAC: 1.30; UGC: 1.22; UUA: 0.26; UCA: 0.97; UAA: 1.36; UGA: 0.80; UUG: 0.92; UCG: 0.80; UAG: 0.85; UGG: 1.00; CUU: 1.45; CCU: 1.48; CAU: 0.87; CGU: 1.16; CUC: 1.77; CCC: 1.19; CAC: 1.13; CGC: 1.44; CUA: 0.52; CCA: 0.86; CAA: 0.88; CGA: 1.59; CUG: 1.08; CCG: 0.47; CAG: 1.12; CGG: 0.47; AUU: 1.13; ACU: 1.09; AAU: 0.60; AGU: 0.70; AUC: 1.59; ACC: 1.27; AAC: 1.40; AGC: 1.17; AUA: 0.29; ACA: 1.07; AAA: 0.47; AGA: 0.78; AUG: 1.00; ACG: 0.56; AAG: 1.53; AGG: 0.55; GUU: 1.31; GCU: 1.40; GAU: 0.98; GGU: 1.36; GUC: 1.59; GCC: 1.34; GAC: 1.02; GGC: 1.43; GUA: 0.39; GCA: 0.74; GAA: 0.75; GGA: 0.92; GUG: 0.71; GCG: 0.52; GAG: 1.25; GGG: 0.30
