## Additional File 9 for "CRISPR/Cas9 mediated editing of the Quorn fungus *Fusarium venenatum* A3/5 by transient expression of Cas9 and sgRNAs targeting endogenous marker gene *PKS12*"

**Table S10 Method for USER cloning of promoter-sgRNA cassettes into AMA1 vectors**

*PCR amplification of inserts*

In a total volume of 50 µl:

1X Phusion™ High-Fidelity DNA Polymerase Buffer

200 µM dNTPs

0.5 µM each primer (F & R)

2 pg template

Phusion™ High-Fidelity DNA Polymerase (Thermo Fisher Scientific) 0.5 units.

PCR cycle

98 °C 30s x1 cycle

98 °C 10s, ‘x’ °C 30s

72 °C 15s per Kb x35 cycles

72 °C 5 min x1 cycle

PCR products were gel purified, eluted in water and quantified by Qubit™ assay

*USER reactions*

Final concentrations of components in a total volume of 12 µl:

50 ng vector linearised and gel purified.

1.5 mM MgCl_2_;

1X Applied Biosystems™ PCR Buffer II without MgCl_2_

USER® enzyme 1.2 units

Purified inserts (eluted in water): for PolII/ribozyme cassettes use equimolar insert:vector ratio;

for 5SRNA cassettes use 5:1 molar insert:vector ratio

Water to a final volume of 12 µl

Incubate at 37 °C 25 minutes, 25 °C 25 minutes

*Recovery of recombinant vectors*

Add entire reaction to 50 µl chemically competent E. coli. Follow protocol provided with cells
