## Additional File 10 for "CRISPR/Cas9 mediated editing of the Quorn fungus *Fusarium venenatum* A3/5 by transient expression of Cas9 and sgRNAs targeting endogenous marker gene *PKS12*"

**Table S11 Method used for protoplast isolation and transformation**

Based on a method described in Moradi *et al* A modified method for transformation of Fusarium graminearum. J. Crop Prot. 2013, 2 (3): 297-304

*Protoplast isolation*

Inoculate 250 ml CMC medium in a 500 ml Erlenmeyer flask with 5 x 4 mm plugs from agar plate culture and incubate for up to 6 days on a rotary shaker at 25°C, shaking at 175 rpm (or to give good aeration). This should yield sufficient protoplasts for up to 40 100 µl aliquots for use in transformation. Filter the culture through 2 layers of sterile Miracloth (Millipore) and centrifuge the filtrate at 4,000×g for 10 min. Discard the supernatant and re-suspend conidia. Place conidia in 100 ml of YEPD broth in a 250 ml Erlenmeyer flask and incubate overnight (14h) in a rotary shaker at 22^o^C, shaking at 175 rpm. Filter the culture through 2 layers of sterile Miracloth using a large sterile funnel and collect the mycelial mat. Rinse the mat with sterile pure water (for consistency, use a volume equal to the volume of the YEPD culture, e.g. 100 ml) and allow the liquid to completely drain by very gently filtering it away using sterile filter papers sandwiched between absorbent tissue. Gently remove the mycelial mat from the miracloth, being careful not to compact it. Place the mycelium into a clean sterile beaker. Add 20 ml Protoplasting Buffer to the beaker and gently disperse the mycelium to ensure efficient digestion. Cover the beaker with sterile foil. Digest for 2 h on a rotary shaker at 28°C, 80 rpm. The digestion time should not be more than 2.5 hours. Filter the digestion mixture through sterile Miracloth (3 layers) into 50 ml sterile conical bottomed plastic tubes. The filtrate should be turbid due to the presence of protoplasts. Centrifuge at RT at 3,000×g for 5 min. Protoplasts are very fragile so treat them gently. Discard the supernatant and gently re-suspend protoplasts in 30 ml of STC Buffer: blow air via a 10 or 25 ml sterile filtered serological pipette and use this and the pipette to gently dislodge the pellet. Centrifuge at 3,000×g for 5 min. Repeat the wash, removing an aliquot for protoplast quantification. Discard the supernatant and gently re-suspend protoplasts in STC Buffer - the amount depends on the number of protoplasts and the final concentration wanted, e.g. from a spore culture of 250 ml re-suspending in up to 4 ml is likely to give approximately 1x10^8^ protoplasts/ml.

It is best to use freshly prepared protoplasts, as their viability declines on freezing and during storage. For storage at ^-^80^o^C first add 40% PEG 4000 in STC (PTC) to give 20% v/v and DMSO to give 7% (e.g. to 2 ml protoplasts in STC, add 500 µl PTC and 175 µl DMSO), mix gently by slow pipetting and dispense 100 µl aliquots into sterile 2 ml tubes and flash freeze in liquid nitrogen before storage at ^-^80^o^C

*Protoplast transformation steps*

- Thaw frozen 100 µl aliquots on ice or use freshly isolated protoplasts prepared as above, held on ice
- Add the DNA (1-5 µg). Flick the tube gently 3 times to mix
- Incubate at RT for 20 min
- Add 1 ml 40% PTC, invert gently three times to mix, incubate at RT for 20 min
- Decant into a 50 ml sterile tube containing 5 ml TB3 supplemented with 100 µg ml^-1^ Hygromycin B, incubate at 21^0^C, shaking at 90 rpm for 14-16 hours (overnight)
- Pour samples into 9 cm petri dishes and add 10 ml Top Agar (melted, cooled to ‘hand hot’ and kept molten in a 50 ^0^C water bath), supplemented with Hygromycin B to give 100 µg ml^-1^ (include one plate with no selection for a non-transformed protoplast control), swirling gently to mix
- Incubate sealed plates for 8-10h at RT, add 10 ml Top Agar prepared as before (supplemented with 100 µg ml ^-1^ Hygromycin B except for the control), seal the plates and incubate at 20^0^C
