## Additional File 11 for "CRISPR/Cas9 mediated editing of the Quorn fungus *Fusarium venenatum* A3/5 by transient expression of Cas9 and sgRNAs targeting endogenous marker gene *PKS12*"

**Table S12 Media used for making protoplasts and protoplast transformation**

| **Carboxymethyl cellulose medium (CMC)** | 1 litre |  | | |  | | | | Stock solution |
| --- | --- | --- | --- | --- | --- | --- | --- | --- | --- |
| Carboxymethyl cellulose sodium salt | 15 | g | | | ***** | | | |  |
| NH_4_ NO_3_ | 1 | g | | |  | | | |  |
| KH_2_ PO_4_ | 1 | g | | |  | | | |  |
| MgSO_4_ ⋅ 7H_2_ O | 0.5 | g | | |  | | | |  |
| Yeast extract | 1 | g | | |  | | | |  |
| *Add slowly to hot water while stirring, continue to heat & stir (dissolves very slowly). Autoclave | | | | | | | | | |
| **YEPD** | 1 litre |  | | |  | | | |  |
| Yeast extract (Oxoid™ LP0021) | 3 | g | | |  | | | |  |
| Peptone (Oxoid™ Bacteriological LP0037) | 10 | g | | |  | | | |  |
| D-glucose | 20 | g | | |  | | | |  |
| 100 ml aliquots in 250 ml flasks. autoclave |  |  | | |  | | | |  |
| **STC buffer** | 1 litre |  | | |  | | | |  |
| Sorbitol 1.2 M | 218.6 | g | | |  | | | | |
| Tris-HCl pH 8.0 10 mM | 10 | ml | | |  | | | | 1M |
| CaCl2 50 mM | 5.5 | g | | |  | | | |  |
| Autoclave |  |  | | |  | | | |  |
| **40% PTC** | 100 ml |  | | |  | | | |  |
| PEG 4000 | 40 | g | | |  | | | |  |
| Dissolve in STC and bring volume to 100 ml. Warm at 65 °C to help dissolve, filter sterilize using a Corning® Bottle-Top Vacuum Filter System with a 0.22 µm nylon membrane. Store at 4 °C | | | | | | | | | |
| **Protoplasting buffer** | 20 ml | | | | | | | | |
| Add to 1.2 M sterile KCl: |  | |  | | | | |  | |
| Driselase (Sigma D9515) | 500 | | mg | | | | |  | |
| Chitinase (sigma C6137) | 1 | | mg | | | | |  | |
| Lysing enzyme (Sigma L1412) | 100 | | mg | | | | |  | |
| NB The Driselase is less soluble, add first and stir for 30-40 minutes then add the others. Make in a sterile beaker using ethanol treated or heat sterilised implements. Filter through a 0.45-mm PES filter | | | | | | | | | |
| **1.2 M KCl** | 1 litre | | |  | |  | | | |
| FW=74.55 | 89.46 | | | g | |  | | | |
| **TB3/Top agar**  Glucose and sorbitol are used as sucrose inhibits hygromycin selection. Make 100 ml aliquots and autoclave | 1 litre | | |  | | |  | | |
| Yeast extract (Oxoid™ LP0021) | 3 | | | g | | |  | | |
| Acid hydrolysed casein | 3 | | | g | | |  | | |
| D-glucose | 10 | | | g | | |  | | |
| Sorbitol | 182 | | | g | | |  | | |
| Agar (omit for TB3) | 15 | | | g | | |  | | |
