## Additional File 12 for "CRISPR/Cas9 mediated editing of the Quorn fungus *Fusarium venenatum* A3/5 by transient expression of Cas9 and sgRNAs targeting endogenous marker gene *PKS12*"

**Table S13: Method for making Potato Sucrose Agar (PSA)**

Ingredients	1 litre

Potato (main crop)	200 g

Sucrose	20 g

Purified agar	20 g

*Method*

- Peel and dice the potatoes (approximately 15-20 mm^3^)
- Boil for 10 mins then filter through gauze
- Add the sucrose to the filtrate, stir to dissolve and make to litre using purified water
- Allow to cool
- Adjust the pH to 6.5 using NaOH
- Aliquot if required
- Add the agar and stir/shake to suspend
- Autoclave to sterilise
