## Additional File 1 for "CRISPR/Cas9 mediated editing of the Quorn fungus *Fusarium venenatum* A3/5 by transient expression of Cas9 and sgRNAs targeting endogenous marker gene *PKS12*"

**Table S1 Expression cassette *Ptef1-FvCas9-SV40-Ttef1* used in this study**

CGAGACAGCAGAATCACCGCCCAAGTTAAGCCTTTGTGCTGATCATGCTCTCGAACGGGCCAAGTTCGGGAAAAGCAAAGGAGCGTTTAGTGAGGGGCAATTTGACTCACCTCCCAGGCAACAGATGAGGGGGGCAAAAAGAAAGAAATTTTCGTGAGTCAATATGGATTCCGAGCATCATTTTCTTGCGGTCTATCTTGCTACGTATGTTGATCTTGACGCTGTGGATCAAGCAACGCCACTCGCTCGCTCCATCGCAGGCTGGTCGCAGACAAATTAAAAGGCGGCAAACTCGTACAGCCGCGGGGTTGTCCGCTGCAAAGTACAGAGTGATAAAAGCCGCCATGCGACCATCAACGCGTTGATGCCCAGCTTTTTCGATCCGAGAATCCACCGTAGAGGCGATAGCAAGTAAAGAAAAGCTAAACAAAAAAAAATTTCTGCCCCTAAGCCATGAAAACGAGATGGGGTGGAGCAGAACCAAGGAAAGAGTCGCGCTGGGCTGCCGTTCCGGAAGGTGTTGTAAAGGCTCGACGCCCAAGGTGGGAGTCTAGGAGAAGAATTTGCATCGGGAGTGGGGCGGGTTACCCCTCCATATCCAATGACAGATATCTACCAGCCAAGGGTTTGAGCCCGCCCGCTTAGTCGTCGTCCTCGCTTGCCCCTCCATAAAAGGATTTCCCCTCCCCCTCCCACAAAATTTTCTTTCCCTTCCTCTCCTTGTCCGCTTCAGTACGTATATCTTCCCTTCCCTCGCTTCTCTCCTCCATCCTTCTTTCATCCATCTCCTGCTAACTTCTCTGCTCAGCACCTCTACGCATTACTAGCCGTAGTATCTGAGCACTTCTCCCTTTTATATTCCACAAAACATAACACAACCTTCACCATGGACAAGAAGTACTCTATCGGCCTCGACATCGGCACCAACTCTGTCGGCTGGGCTGTCATCACCGACGAGTACAAGGTCCCTTCTAAGAAGTTCAAGGTCCTCGGCAACACCGACCGACACTCTATCAAGAAGAACCTCATCGGCGCTCTCCTCTTCGACTCTGGCGAaACCGCTGAGGCTACCCGACTCAAGCGAACCGCTCGACGACGATACACCCGACGAAAGAACCGAATCTGCTACCTCCAGGAGATCTTCTCTAACGAGATGGCTAAGGTCGACGACTCTTTCTTCCACCGACTCGAGGAGTCTTTCCTCGTCGAGGAGGACAAGAAGCACGAGCGACACCCTATCTTCGGCAACATCGTCGACGAGGTCGCTTACCACGAGAAGTACCCTACCATCTACCACCTCCGAAAGAAGCTCGTCGACTCTACCGACAAGGCTGACCTCCGACTCATCTACCTCGCTCTCGCTCACATGATCAAGTTCCGAGGCCACTTCCTCATCGAGGGCGACCTCAACCCTGACAACTCTGACGTCGACAAGCTCTTCATCCAGCTCGTCCAGACCTACAACCAGCTCTTCGAGGAGAACCCTATCAACGCTTCTGGCGTCGACGCTAAGGCTATCCTCTCTGCTCGACTCTCTAAGTCTCGACGACTCGAGAACCTCATCGCTCAGCTCCCTGGCGAGAAGAAGAACGGCCTCTTCGGCAACCTCATCGCTCTCTCTCTCGGCCTCACCCCTAACTTCAAGTCTAACTTCGACCTCGCTGAGGACGCTAAGCTCCAGCTCTCTAAGGACACCTACGACGACGACCTCGACAACCTCCTCGCTCAGATCGGCGACCAGTACGCTGACCTCTTCCTCGCTGCTAAGAACCTCTCTGACGCTATCCTCCTCTCTGACATCCTCCGAGTCAACACCGAGATCACCAAGGCTCCTCTCTCTGCTTCTATGATCAAGCGATACGACGAGCACCACCAGGACCTCACCCTCCTCAAGGCTCTCGTCCGACAGCAGCTCCCTGAGAAGTACAAGGAGATCTTCTTCGACCAGTCTAAGAACGGCTACGCTGGCTACATCGACGGCGGCGCTTCTCAGGAGGAGTTCTACAAGTTCATCAAGCCTATCCTCGAGAAGATGGACGGCACCGAGGAGCTCCTCGTCAAGCTCAACCGAGAGGACCTCCTCCGAAAGCAGCGAACCTTCGACAACGGCTCTATCCCTCACCAGATCCACCTCGGCGAGCTCCACGCTATCCTCCGACGACAGGAGGACTTCTACCCTTTCCTCAAGGACAACCGAGAGAAGATCGAGAAGATCCTCACCTTCCGAATCCCTTACTACGTCGGCCCTCTCGCTCGAGGCAACTCTCGATTCGCTTGGATGACCCGAAAGTCTGAGGAaACCATCACCCCTTGGAACTTCGAGGAGGTCGTCGACAAGGGCGCTTCTGCTCAGTCTTTCATCGAGCGAATGACCAACTTCGACAAGAACCTCCCTAACGAGAAGGTCCTCCCTAAGCACTCTCTCCTCTACGAGTACTTCACCGTCTACAACGAGCTCACCAAGGTCAAGTACGTCACCGAGGGCATGCGAAAGCCTGCTTTCCTCTCTGGCGAGCAGAAGAAGGCTATCGTCGACCTCCTCTTCAAGACCAACCGAAAGGTCACCGTCAAGCAGCTCAAGGAGGACTACTTCAAGAAGATCGAGTGCTTCGACTCTGTCGAGATCTCTGGCGTCGAGGACCGATTCAACGCTTCTCTCGGCACCTACCACGACCTCCTCAAGATCATCAAGGACAAGGACTTCCTCGACAACGAGGAGAACGAGGACATCCTCGAGGACATCGTCCTCACCCTCACCCTCTTCGAGGACCGAGAGATGATCGAGGAGCGACTCAAGACCTACGCTCACCTCTTCGACGACAAGGTCATGAAGCAGCTCAAGCGACGACGATACACCGGCTGGGGCCGACTCTCTCGAAAGCTCATCAACGGCATCCGAGACAAGCAGTCTGGCAAGACCATCCTCGACTTCCTCAAGTCTGACGGCTTCGCTAACCGAAACTTCATGCAGCTCATCCACGACGACTCTCTCACCTTCAAGGAGGACATCCAGAAGGCTCAGGTtTCTGGCCAGGGCGACTCTCTCCACGAGCACATCGCTAACCTCGCTGGCTCTCCTGCTATCAAGAAGGGCATCCTCCAGACCGTCAAGGTCGTCGACGAGCTCGTCAAGGTCATGGGCCGACACAAGCCTGAGAACATCGTCATCGAGATGGCTCGAGAGAACCAGACCACCCAGAAGGGCCAGAAGAACTCTCGAGAGCGAATGAAGCGAATCGAGGAGGGCATCAAGGAGCTCGGCTCTCAGATCCTCAAGGAGCACCCTGTCGAGAACACCCAGCTCCAGAACGAGAAGCTCTACCTCTACTACCTCCAGAACGGCCGAGACATGTACGTCGACCAGGAGCTCGACATCAACCGACTCTCTGACTACGACGTCGACCACATCGTCCCTCAGTCTTTCCTCAAGGACGACTCTATCGACAACAAGGTCCTCACCCGATCTGACAAGAACCGAGGCAAGTCTGACAACGTCCCTTCTGAGGAGGTCGTCAAGAAGATGAAGAACTACTGGCGACAGCTCCTCAACGCTAAGCTCATCACCCAGCGAAAGTTCGACAACCTCACCAAGGCTGAGCGAGGCGGCCTCTCTGAGCTCGACAAGGCTGGCTTCATCAAGCGACAGCTCGTCGAaACCCGACAGATCACCAAGCACGTCGCTCAGATCCTCGACTCTCGAATGAACACCAAGTACGACGAGAACGACAAGCTCATCCGAGAGGTCAAGGTCATCACCCTCAAGTCTAAGCTCGTtTCTGACTTCCGAAAGGACTTCCAGTTCTACAAGGTCCGAGAGATCAACAACTACCACCACGCTCACGACGCTTACCTCAACGCTGTCGTCGGCACCGCTCTCATCAAGAAGTACCCTAAGCTCGAGTCTGAGTTCGTCTACGGCGACTACAAGGTCTACGACGTCCGAAAGATGATCGCTAAGTCTGAGCAGGAGATCGGCAAGGCTACCGCTAAGTACTTCTTCTACTCTAACATCATGAACTTCTTCAAGACCGAGATCACCCTCGCTAACGGCGAGATCCGAAAGCGACCTCTCATCGAaACCAACGGCGAaACCGGCGAGATCGTCTGGGACAAGGGCCGAGACTTCGCTACCGTCCGAAAGGTCCTCTCTATGCCTCAGGTCAACATCGTCAAGAAaACCGAGGTCCAGACCGGCGGCTTCTCTAAGGAGTCTATCCTCCCTAAGCGAAACTCTGACAAGCTCATCGCTCGAAAGAAGGACTGGGACCCTAAGAAGTACGGCGGCTTCGACTCTCCTACCGTCGCTTACTCTGTCCTCGTCGTCGCTAAGGTCGAGAAGGGCAAGTCTAAGAAGCTCAAGTCTGTCAAGGAGCTCCTCGGCATCACCATCATGGAGCGATCTTCTTTCGAGAAGAACCCTATCGACTTCCTCGAGGCTAAGGGCTACAAGGAGGTCAAGAAGGACCTCATCATCAAGCTCCCTAAGTACTCTCTCTTCGAGCTCGAGAACGGCCGAAAGCGAATGCTCGCTTCTGCTGGCGAGCTCCAGAAGGGCAACGAGCTCGCTCTCCCTTCTAAGTACGTCAACTTCCTCTACCTCGCTTCTCACTACGAGAAGCTCAAGGGCTCTCCTGAGGACAACGAGCAGAAGCAGCTCTTCGTCGAGCAGCACAAGCACTACCTCGACGAGATCATCGAGCAGATCTCTGAGTTCTCTAAGCGAGTCATCCTCGCTGACGCTAACCTCGACAAGGTCCTCTCTGCTTACAACAAGCACCGAGACAAGCCTATCCGAGAGCAGGCTGAGAACATCATCCACCTCTTCACCCTCACCAACCTCGGCGCTCCTGCTGCTTTCAAGTACTTCGACACCACCATCGACCGAAAGCGATACACCTCTACCAAGGAGGTCCTCGACGCTACCCTCATCCACCAGTCTATCACCGGCCTCTACGAaACCCGAATCGACCTCTCTCAGCTCGGCGGCGACTGA***TG*CCTCCCAAGAAGAAGCGCAAGGTCTGAGCGGACATTCGATTTATGCCGTTATGACTTCCTTAAAAAAGCCTTTACGAATGAAAGAAATGGAATTAGACTTGTTATGTAGTTGATTCTACAATGGATTATGATTCCTGAACTTCAAATCCGCTGTTCATTATTAATCTCAGCTCTTCCCGTAAAGCCAATGTTGAAACTATTCGTAAATGTACCTCGTTTTGCGTGTACCTTGCTTATCACGTGATATTACATGACCTGGACAGAGTTCTGCGCGAAAGTCATAACGTAAATCCCGGGCGGTAGGTGCGTCCCGGGCGGAAGGTAGTTTTCTCGTCCACCCCAACGCGTTTATCAACCTCAACTTTCAACAACCATCATGCCACCAAAAGCGCGTAAAACAAAGCGAGATTTGATTGAGCAAGAGGGCAGGATCCAATGCGCGATTCAAGACATTAAAAATGGAAAATTTCAAAAAATTGCGCCCGCAGCGCGTGCATACAAAATTCATCCCAATAC**

Constituent parts of the *Ptef1-FvCas9-SV40-Ttef1* cassette are denoted by different font colours: red font is sequence for *Ptef1*; Black font is coding sequence for the *Cas9* gene codon-optimised for Fusarium venenatum (*Fv_Cas9*) used in this study; pink bold font is sequence for SV40_NLS_-STOP codon (preceding bases in italics were added for frameshift) ; blue bold font is sequence for *Ttef1*.
