## Additional File 2 for "CRISPR/Cas9 mediated editing of the Quorn fungus *Fusarium venenatum* A3/5 by transient expression of Cas9 and sgRNAs targeting endogenous marker gene *PKS12*"

**Table S2 Sequence of *P5SrRNA*-sgRNA cassettes used in this study**

*P5SrRNA* sgRNA cassettes CACATACGACCAAAGGTAGTGGAAAATACGGGATCCCGTCCGCTCTCCCATAGTCAAGCCACTAACCGGCGGATTAGTAGTTGGGTCGGTGACGACCAGCGAATCCCCGCTGTTGTATGTNNNNNNNNNNNNNNNNNNNNGTTTTAGAGCTAGAAATAGCAAGTTAAAATAAGGCTAGTCCGTTATCAACTTGAAAAAGTGGCACCGAGTCGGTGC**TTTTTT***GTAGTAACACCCGGAGTGCATGGATCA*

Capital font = 5SRNA promoter sequence; ‘N’s = *PKS12* gene target site sequences (PK3-14 CCAGAACAGCCTGAAATGAG; PK3-29 CACCATAACATGACCATTAG); underlined capital font = scaffold sequence; bold font = terminator sequence; capital italics = spacer sequence
